# Old and ancient trees are life history lottery winners and act as evolutionary buffers against long-term environmental change

**DOI:** 10.1101/2021.10.16.464670

**Authors:** Charles H. Cannon, Gianluca Piovesan, Sergi Munné-Bosch

## Abstract

Trees can live many centuries with sustained fecundity and death is largely stochastic. We use a neutral stochastic model to examine the demographic patterns that emerge over time, across a range of population sizes and empirically observed mortality rates. A small proportion of trees (∼1% at 1.5% mortality) are life-history ‘lottery’ winners, achieving ages >10-20x median age. Maximum age increases with bigger populations and lower mortality rates. One quarter of trees (∼24%) achieve ages that are 3-4 times greater than median age. Three age classes (Mature, Old, and Ancient) contribute unique historical diversity across complex environmental cycles. Ancient trees are an emergent property of forests that requires many centuries to generate. They rradically change generation time variance and population fitness, bridging infrequent environmental cycles. These life-history ‘lottery’ winners are vital to future forest dynamics and invaluable data about environmental history and individual longevity. Old-growth forests contain trees that cannot be replaced through restoration or regeneration in the near future. They simply must be protected to preserve their unique diversity.

## Old Trees in Nature

Human cultures around the world have revered ancient trees as powerful spiritual beings, connecting Earth and Heaven, as sources of wisdom, fertility, balance, and longevity ^1^. These myths and legends are often embodied by a particularly old and venerated individual tree of exceptional presence, distinguishable from the many other large trees in the forest. These ancient trees are seen as belonging to a separate and special class of being that transcends the normal plane of existence, binding the tree to a deep knowledge and awareness of history, change, and persistence. With increasing knowledge about the role old trees play in ecosystems, biologists are also beginning to attach special significance to individual ancient trees in populations ^2, 3^.

The ecological importance of old trees in forested ecosystems has been extensively documented, particularly as small natural features that provide a wide range of services ^4–6^. And yet, our understanding of tree age structure in forested ecosystems remains poor ^7^. Fundamentally, the lifespan of even an ordinary tree exceeds the duration of long-term ecological projects, so demographic studies of trees can only be performed on a cross-sectional, not longitudinal, basis. Additionally, several inherent difficulties exist for the reliable dating of old trees ^8, 9^. Among tropical tree species in aseasonal climates, growth rings are not tightly linked to annual cycles, making temporal dynamics difficult to interpret ^10-11^. Overall, the most reliable methods require complex – time consuming - studies and these are rare at the population or community level ^12^.

Annual mortality rates have been estimated and measured at the population or community level in many forests. These rates range from 0.3% to 5% for large mature well-established individuals ^13–19^, often centered around 1.5-2%. While individual-based stochastic models of tree mortality are commonly used to explore community ecological and structural dynamics ^7^, the evolutionary implications of extreme longevity of individuals have not previously been noted or explored. Using a neutral stochastic mortality model, we obtain results that closely match those obtained through empirical studies ^20–22^. Here, we highlight several exceptional population-level demographic properties that emerges from a neutral mortality model based upon empirical studies and discuss their evolutionary implications, particularly given long-term environmental cycles.

First, a very small proportion of individuals win the longevity “lottery” and obtain exceptional ages, >>10 times greater than the median age in the population (Figure 1A). These ancient trees are observed in natural populations and are possible because of the lack of programmed senescence enabled by the woody plant growth form ^3^ and the low mortality rates observed in many old growth forests globally ^13–16^. We argue that despite the rarity of these individuals, they play a significant role maintaining diversity in the population and bridging across unusual and infrequent environmental conditions. Second, a larger proportion of individuals reach significant ages many times greater than the median age. These old individuals contribute substantially to the stabilization of population diversity to intermediate environmental change. We objectively identify three age classes (“mature”, “old”, and “ancient”, see Methods for classification technique), examine the properties of these groups in response to variation in population size and mortality rate, and explore how each group relates to both short-term and long-term environmental changes.

**Figure 1.**
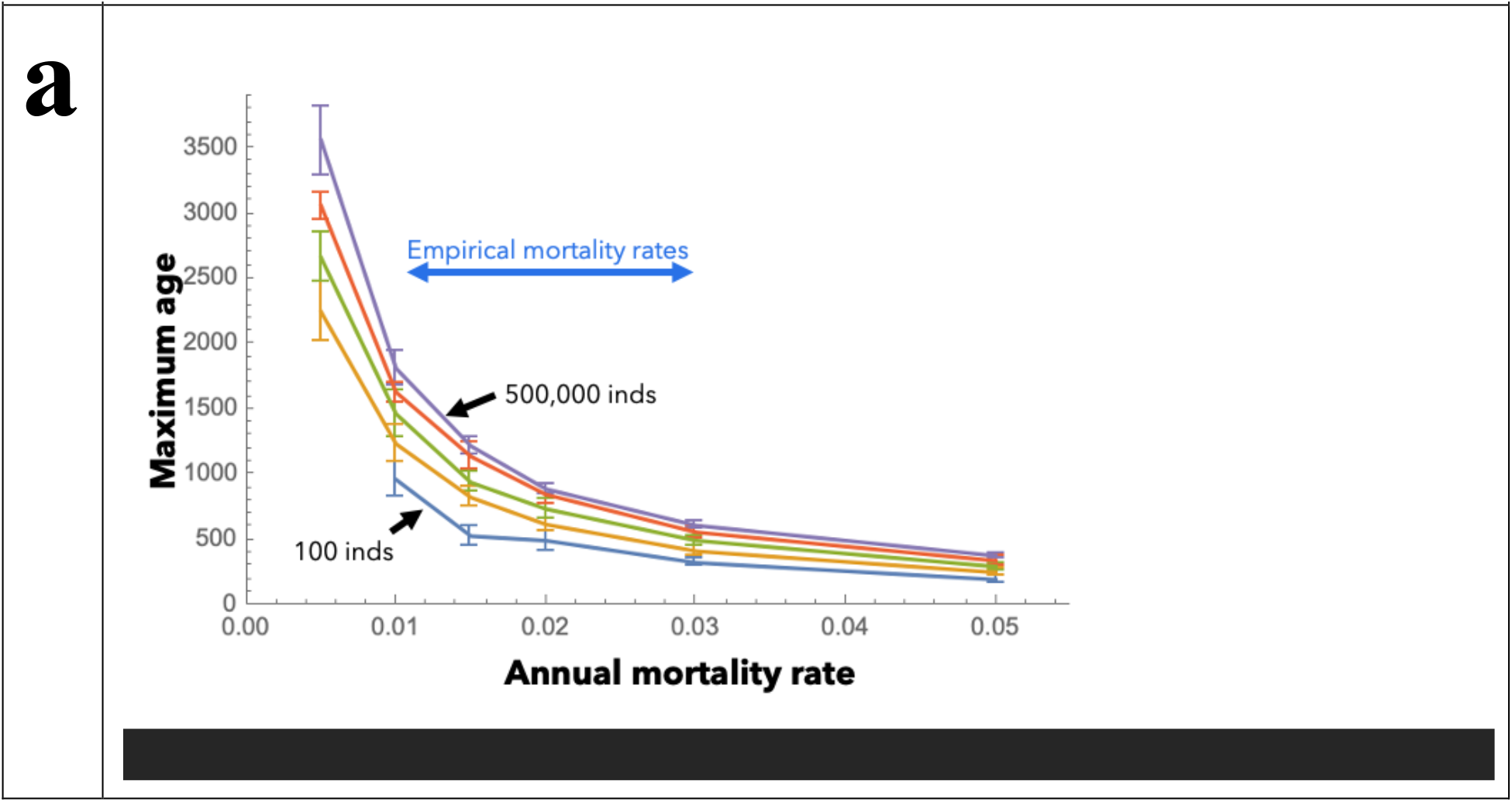
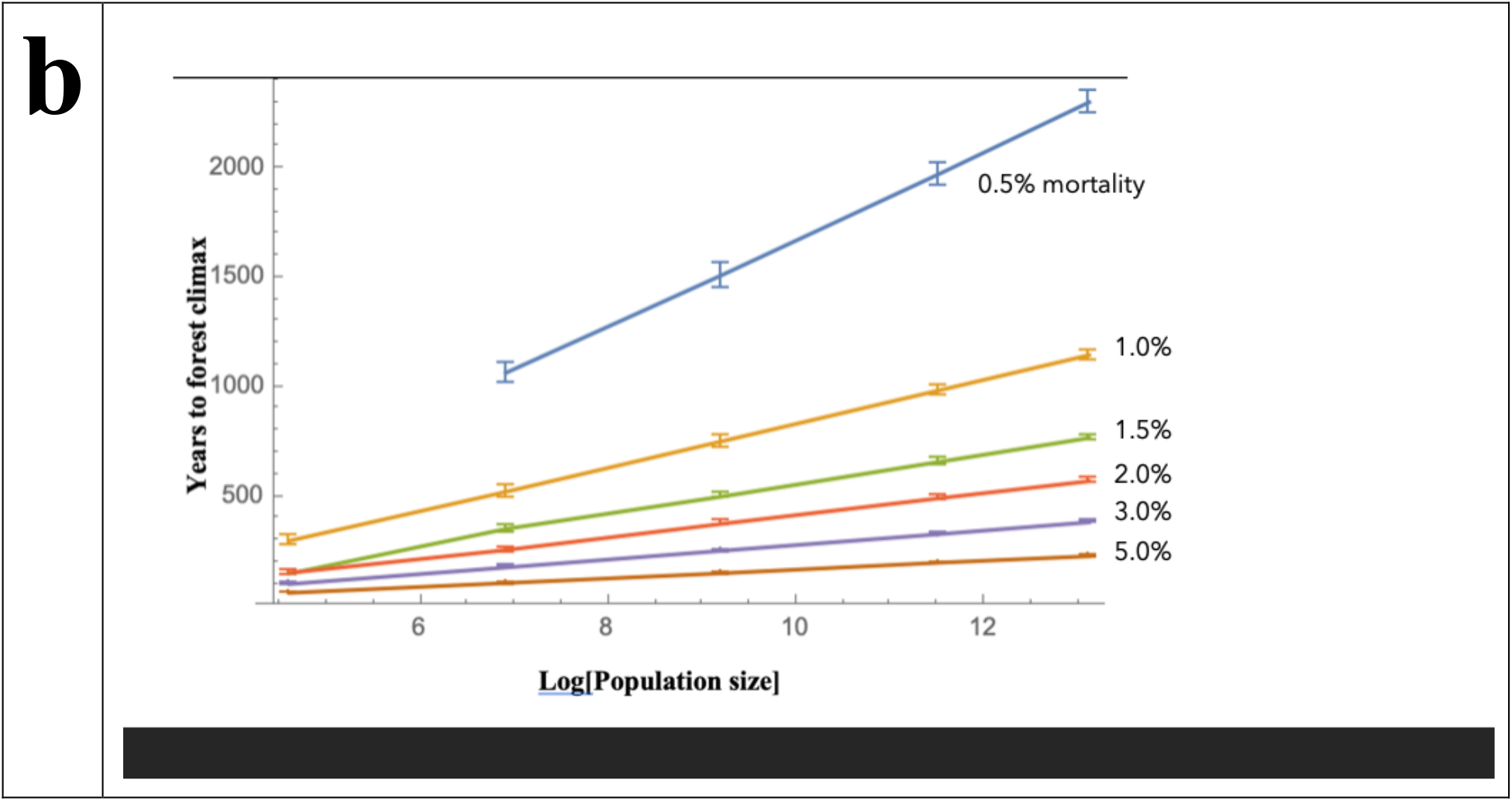
Patterns of individual tree and forest age, given different population sizes and annual stochastic mortality rates. **A)** Maximum age of trees, given different population size (colored lines) and mortality rate (x-axis), across 25 replicates for each parameter set. Each line indicates steady state population size during each replicate, which decreases top to bottom (purple = 500K; red = 100K; green = 10K; orange = 1K; blue = 100). Annual stochastic mortality rates examined: 0.005, 0.01, 0.015, 0.02, 0.03, 0.05. **B)** Climax age for forests of different size and mortality rates. The population size is shown on the x-axis while each line indicates the forest climax age, given decreasing annual mortality rates, top to bottom (blue = 0.005%; red = 0.01%; green = 0.015%; black = 0.02%; orange = 0.03%; purple = 0.05%). The extension of the green line illustrates the trend going up to one million individuals. Climax age is defined here as the point at which the oldest class of individuals only has one individual, after establishing from a single uniform aged cohort (see text for further explanation).

## Modeling tree evolutionary demographics

Mature, well-established trees are not programmed to senesce at a particular size or age but instead die in consequence of serious damage due to abiotic factors, such as fires or severe storms or sustained poor environmental conditions. In our model, we assume that established canopy trees would survive until stochastic death causes mortality. We are only modeling mature established trees and not seedlings, which are known to experience very high mortality rates. We explore these demographic dynamics across a range of basic parameters for annual mortality rate (0.1, 0.5, 1.0, 1.5, 2.0, 3.0, and 5.0%); and population size (100, 1K, 10K, 100K, and 500K individuals); given a time span of 15K years. Population sizes are held stable throughout the simulation, assuming a mature old-growth forest where space is fully occupied and community size constrained. Population age refers to the length of time after establishment of the first cohort of mature trees. Twenty-five replicates were run for each set of model parameters and results reflect the average among replicates. These simulations are neutral in process to examine the understanding patterns that emerge, given the basic assumptions of the model (see Methods for more detailed description of the process). Code was written in Mathematica 12 (see Supplementary Material).

The existence of life history ‘lottery’ winners is apparent in all model conditions, but the maximum age reached increases dramatically as mortality rate declines below 2% (Fig. 1A), with a doubling of maximum age between 1% and 0.5% mortality. This range of mortality rates corresponds directly to empirical values estimated from global forests, indicating the small changes in forest mortality can have dramatic impact on the potential age of trees in that forest. Additionally, maximum age does respond to increasing population size but this effect declines for very large populations, with the most dramatic response occurring among smaller populations, indicating the sensitivity of small forest fragments to this process. Median population age, on the other hand, is insensitive to population size but declines sharply with increasing mortality rate (Fig. 2). The mean age of old also increases with declining mortality rate and increasing population size, although less dramatically. Interestingly, the proportion of Old Trees in the population (∼24.4%) remains constant across all model conditions while the proportion of Ancients remains constant for a particular mortality rate (e.g. ∼1% of population for 1.5% mortality) but increases with declining mortality (e.g. ∼1.5% of population for 1% mortality compared to 0.2% of population for 5% mortality). The Ancient trees, particularly the oldest, display the greatest degree of responsiveness to changing model conditions.

**Figure 2.**
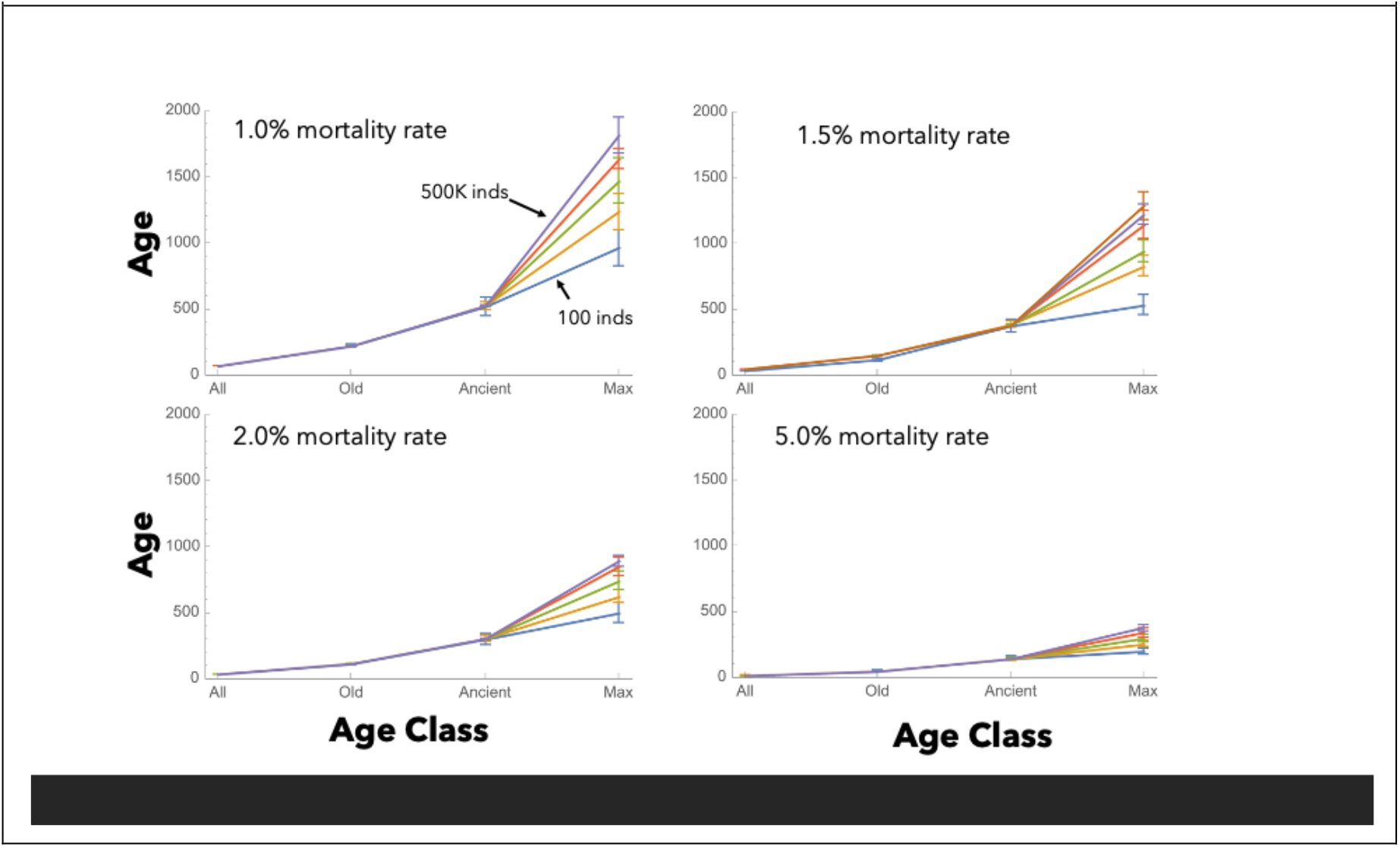
Mean age and standard deviation of different age classes, given different mortality rates and population sizes, across replicates. The lines connect models with different population sizes, with the smallest population size (100 inds) having the lowest maximum up to the largest population size (500K inds) with the highest maximum age.

Our models also provide an objective estimate of the time required for a forest to reach climax age, following the establishment of an even-aged cohort. This situation resembles what happens managed restoration or natural regeneration after catastrophic fire or storm or anthropogenic conversion. The climax age of a forest is a linear relationship between Log[population size] and mortality rate (Fig. 1B), taking several centuries given intermediate population sizes and mortality rates below 2%. We define this climax age as the point where the age structure of the population stabilizes (Supplementary Material 1) and the forest becomes older than the oldest surviving individual. This moment in forest evolution also corresponds closely to when the oldest individual becomes the lone representative of its age. These patterns have relevance to many modern forests, particularly in the northern temperate regions, where even-aged stands have grown in abandoned agricultural lands. Given prevalent natural mortality rates, the climax status of the forest remains centuries in the future. With increased mortality rates observed in forests around the world, these dynamics may be accelerated and the potential for the emergent ancient age groups may be impossible.

We defined threshold ages for three age groups (Mature, Old, and Ancient) based upon the properties of the rank-age distribution of the populations, after the population had reached climax age (Supplementary Material 2). While the mature and old age groups remain consistent across model conditions, we would highlight the variability and sensitivity of the ancient age group to changes in mortality rate. Some individuals obtain truly astonishing ages in relation to the mean age of the oldest individual, which as noted above, is already substantially greater than median population age and even the median oldest individual across replicates (Fig. 3). Even in the smallest populations (Fig. 2), individuals achieve ages that are significantly more than the mean maximum age and the frequency and magnitude of these outliers increase with declining mortality. Essentially, a life-history lottery winner can emerge at any time point. The maximum age obtained by these lottery winners is substantially greater than even forest climax age, indicating that even after population age structure has stabilized, the ancient trees continue to become more different and unusual for many more centuries. Unfortunately, the ancient age group that emerges from a stochastic death process, and thus their impact on evolutionary dynamics, can only be found in old-growth forests. Modern conversion resets the long process.

**Figure 3.**
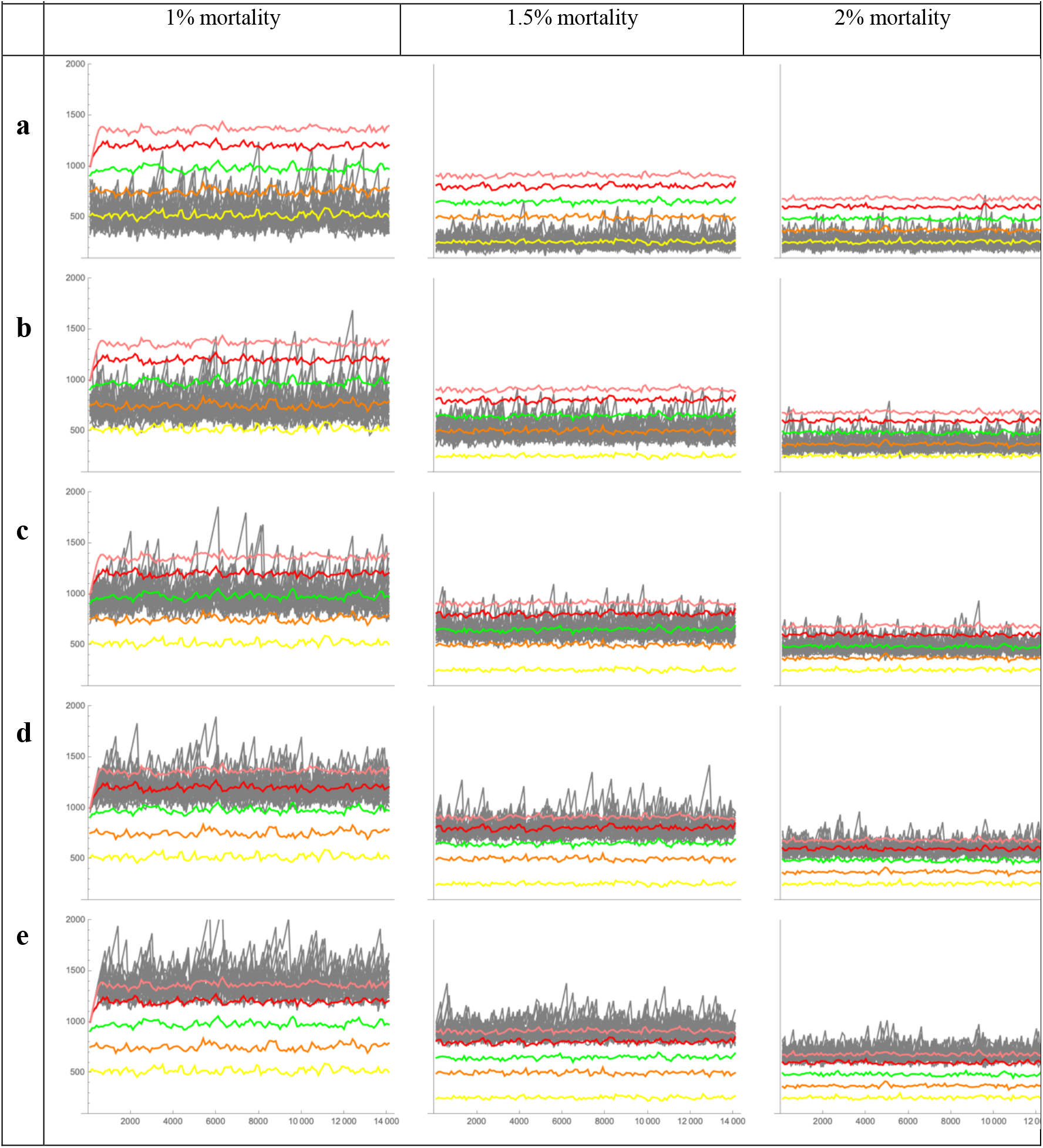
Oldest individual, given different population sizes and annual mortality rates, across 25 replicates over 150K years. In each figure, time is shown on the x-axis, running from 1000 to 150K years and maximum age is show on the y-axis. Different mortality rates are shown in each column of figures. The solid-colored lines in each figure illustratea the average age of the oldest individual at each time point across the 25 reps for each population size: yellow = 100; orange = 1K; green = 10K; red = 100K; burnt orange = 500K, given the corresponding mortality rate. The gray lines in the background illustrate the maximum age in all 25 replicates for each population size, by row: A=100, B=1K, C=10K, D=100K, E=500K).

## Buffering evolutionary change across different time scales

Environmental change occurs on many different time scales and is an emergent selection force that is composed of numerous factors of change. To explore the impact of the three age groups on population level diversity, we constructed a scenario of millennial environmental change, where fitness is determined by four different underlying regular cycles (Fig. 4 and Supplementary Material 3). Given a mortality rate of 1.5%, fitness diversity in each of the three age groups varies with increasing population size in different ways. Compared to the overall distribution of fitness values present in the environment over one thousand years, the Mature individuals correspond extremely well with fitness values over the short-term but poorly with fitness averaged over the entire time period. This distribution for Mature individuals does not change with population size because this age group is present and consistent in all populations. The Old individuals, on the other hand, closely match fitness diversity in the Mature group at the smallest population size (Fig. 4a) but quickly shift until they closely match the distribution of environmental variation. Fitness diversity among Ancient individuals continues to shift with each change in population size. At the smallest population size, Ancient individuals are entirely absent and as they appear, they are substantially different than either of the two age groups and the environmental change. With increasing population size and a corresponding increase in abundance and age of ancient individuals, their fitness diversity begins to converge on the distribution of overall environmental variation (Fig. 4e).

**Figure 4.**
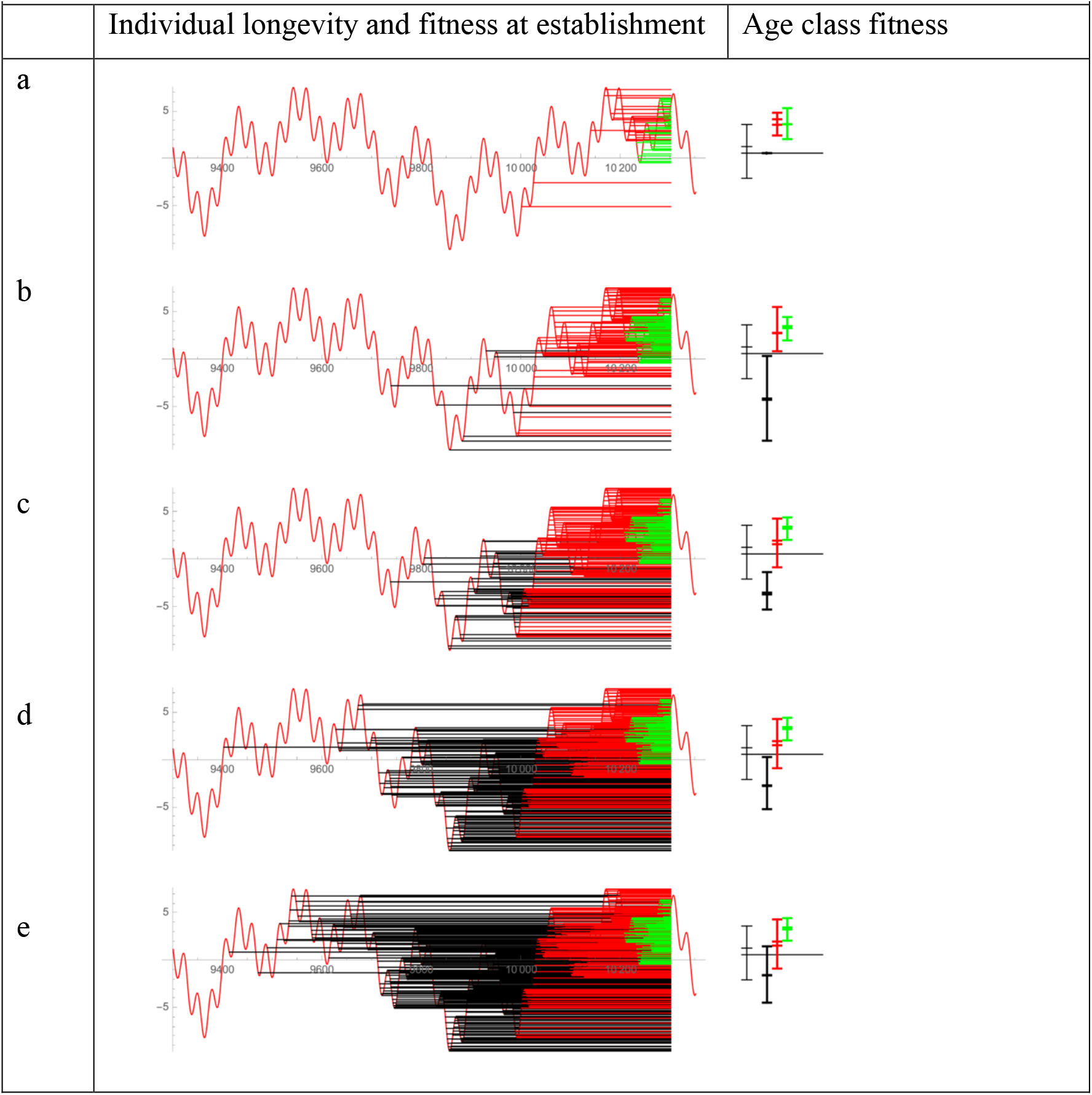
Representative population demographics and fitness through time, given an annual stochastic mortality rate of 1.5% and an environmental fitness that varies through time. Each row of figures corresponds to a different population size: a) 100, b) 1K, c) 10K, d) 100K, e) 500K. In the left column, the individual longevity of trees is shown by a line, running left to right, from the time of their establishment to the time. Lines are color-coded according to age group: green = mature; red = old; black = ancients (see Methods for technical definitions). Each figure represents the population of living individuals observed at a single sample time (10,300 years) from a single replicate. The sample time was chosen for illustration purposes. The environmental fitness value for all is shown by the red waveform populations (y-axis), which is the outcome of four underlying environmental patterns (see Supplemental material) changing through time (x-axis). In the right column, the descriptors of age group fitness are shown. The error bars illustrate the 25% quantile of population fitness, the lower central bar corresponds to population mean, the upper central bar corresponds to population median. The thin black errors bars illustrate the environmental fitness value over the one thousand year period, including the horizontal black line, which corresponds to the mean environmental fitness value over this time period.

The environmental scenario discussed here represents a single set of conditions for the scale and frequency of environmental change and are thus only a case study but certain lessons can be drawn from its results. First, Mature individuals are present and completely predictable in all populations, no matter the model parameters, although median age does decline with increasing mortality rate. These Mature individuals are constrained to environmental conditions experienced in the recent past. Secondly, the Old individuals are also present and consistent in all but the smallest populations and reflect the central tendency of environmental variation, given that the time scale of change and the mean age of Old individuals roughly corresponds. Finally, the Ancient individuals in intermediate sized populations can poorly fit both current and historical environmental variation and represent a unique range of diversity compared to the other age groups.

These Ancient trees can be a very valuable resource for the population as a whole, if temporal scales of environmental variation extend beyond the age of old individuals, thus bridging between extreme and infrequent environmental conditions that the population might not survive without the ancient trees. Conversely, these ancient trees might harbor traits and genes that are poorly fit to current and more short-termed environmental change and if they dominated reproduction could reduce population fitness. This could be particularly concerning if environmental change were directional, moving steadily away from any central tendency. Ultimately, many other possible environmental dynamics could be explored, particularly if environmental change is directional and not constrained to a central value. Their rarity and difficult identification makes discovery and direct study challenging using traditional ecological sampling methods. Instead, the disproportionate and profound impact these ancient trees have on the adaptiveness and sustainability of the population and species as a whole require an innovative and specialized modeling approach. A full exploration of these dynamics requires a comprehensive analytical approach to summarize results across a range of environmental conditions to obtain general properties of this dynamic.

To assess the evolutionary buffering capacity of the emerging age class structure in these populations, we examined variation in three estimators of generation time, given different population sizes and mortality rates. The first estimator assumes that all individuals alive in the population have an equal probability of producing successful offspring. The second estimator assumes that older individuals contribute disproportionately to the next generation, based upon their age and not the number of individuals of that age. The third estimator assumes that only the oldest individual alive produces offspring. The first two estimators are basic methods of estimating generation time while the last value indicates the maximum possible value for generation time in a population over a given period of time.

Generation time for all three estimators increased with decreasing mortality rate, with only slight changes in the first estimator but increased roughly sixfold for the last estimator (Table 2). These values also changed dramatically with increasing population size, increasing two- to three-fold within each mortality rate. The first and third estimators essentially represent the lower and upper bounds for generation time in the population. Given that selective environments change through a complex cyclical process of several underlying patterns, extreme environmental conditions can return over time periods of decades or hundreds of years (Figure 4A). Greater longevity of ancient individuals increasingly bridges the temporal gap between the return of these environmental extremes (Fig. 4B-E). If individuals that establish during those extreme periods are more fit for those conditions, they can produce offspring that are likely to have advantageous alleles that facilitate establishment during those extreme conditions. Because fecundity is generally maintained in ancient trees, their contribution to regeneration during these extreme but rare climatic conditions. Overall, despite the rarity of the ancient individuals, each ancient individual is connected to a unique historical circumstance (Fig. 4). They create a rich and deep genetic diversity within the community that can bridge the gaps between rare and extraordinary environmental conditions. Particularly, given predictions of global climate change, the baseline itself of cyclical environmental change is shifting and the amplitude of change is increasing, driving conditions towards more and more extreme values, further accentuating the importance of these ancient reservoirs of valuable adaptive capacity at the outer margins of genetic diversity. Beyond the neutral dynamics explored here, ancient trees also accumulate heritable epigenetic imprinting and somatic mutations which further increase the overall population diversity.

## Mechanisms to deal with aging in ancient trees

Several mechanisms have evolved in individual trees to enable extreme longevity and deal with negative effects of ageing (Fig. 4). In general, they can be grouped into two non-mutually exclusive categories: (i) senescence avoidance and (ii) aging tolerance. “Senescence avoidance” includes either escaping from or preventing senescence. Clonality in plants allows resetting the clock and escaping the wear and tear of ageing ^23^. Although clonality is tightly linked to long lifespans, it only relates to the genotype, but not the individual ^3,24,25^. More relevant to the model presented here, ancient, non-clonal trees usually show another mechanism to prevent aging for a very significant part of their lifespan. Growth and senescence at two essential parts of the same coin. While plants are growing, they cannot senesce. Senescence implies a reduction in vigour; therefore, continuous growth is the most effective mechanism to prevent senescence. Several long-lived plant species follow this strategy, including redwoods (*Sequoia sempervirens*), which attain their maximum height at relatively advanced ages and generally do not show a decline in vegetative growth vigour with increasing age ^26^. This strategy has, however, its limits. Due to functional leaf characteristics in redwoods, increasing leaf water stress due to hydraulic and mechanical constraints may ultimately limit leaf expansion and photosynthesis, thereby preventing further height growth beyond 130 meters even with ample soil moisture ^27^. Trees have evolved strategies to prevent death from water stress once maximum height has been reached.

Stress tolerance is indeed a mechanism of aging tolerance, since it serves to delay death as trees age. In ancient trees, the ability to maintain pluripotent meristems is key for resistance (growth memory), resilience and stability of populations ^28,29^. While shoot apical meristems (SAMs) lead directly to vegetative and reproductive growth, axillary meristems (AMs) are particularly important for plant branching and regeneration after damage ^30^. Ancient trees possess huge vegetative plasticity (Fig. 5), even through epicormic shoots in consequence of severe disturbances ^3^.

**Figure 5.**
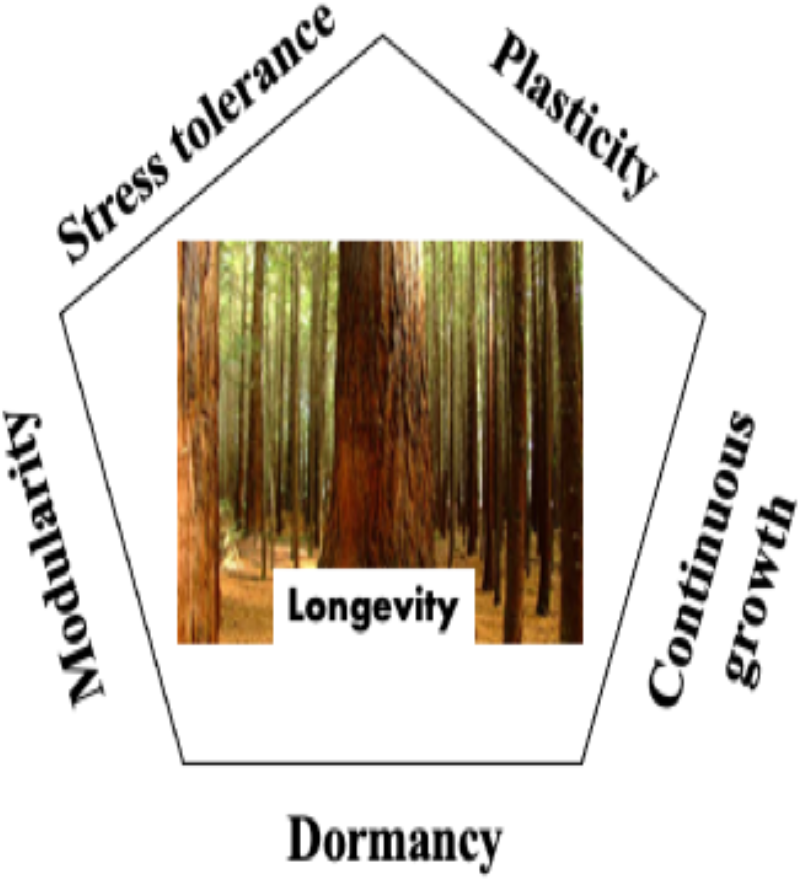
Major mechanisms evolved by ancient trees to defy aging. Lack of programmed senescence enables very long lifespans in trees and implies that death is largely due to stochastic events. The indeterminate life span of trees arise from a combination of mechanisms that serve both to prevent senescence (modularity, continuous growth, dormancy) and tolerate aging (stress tolerance), creating enormous potential and flexibility in longevity.

**Figure 6.**
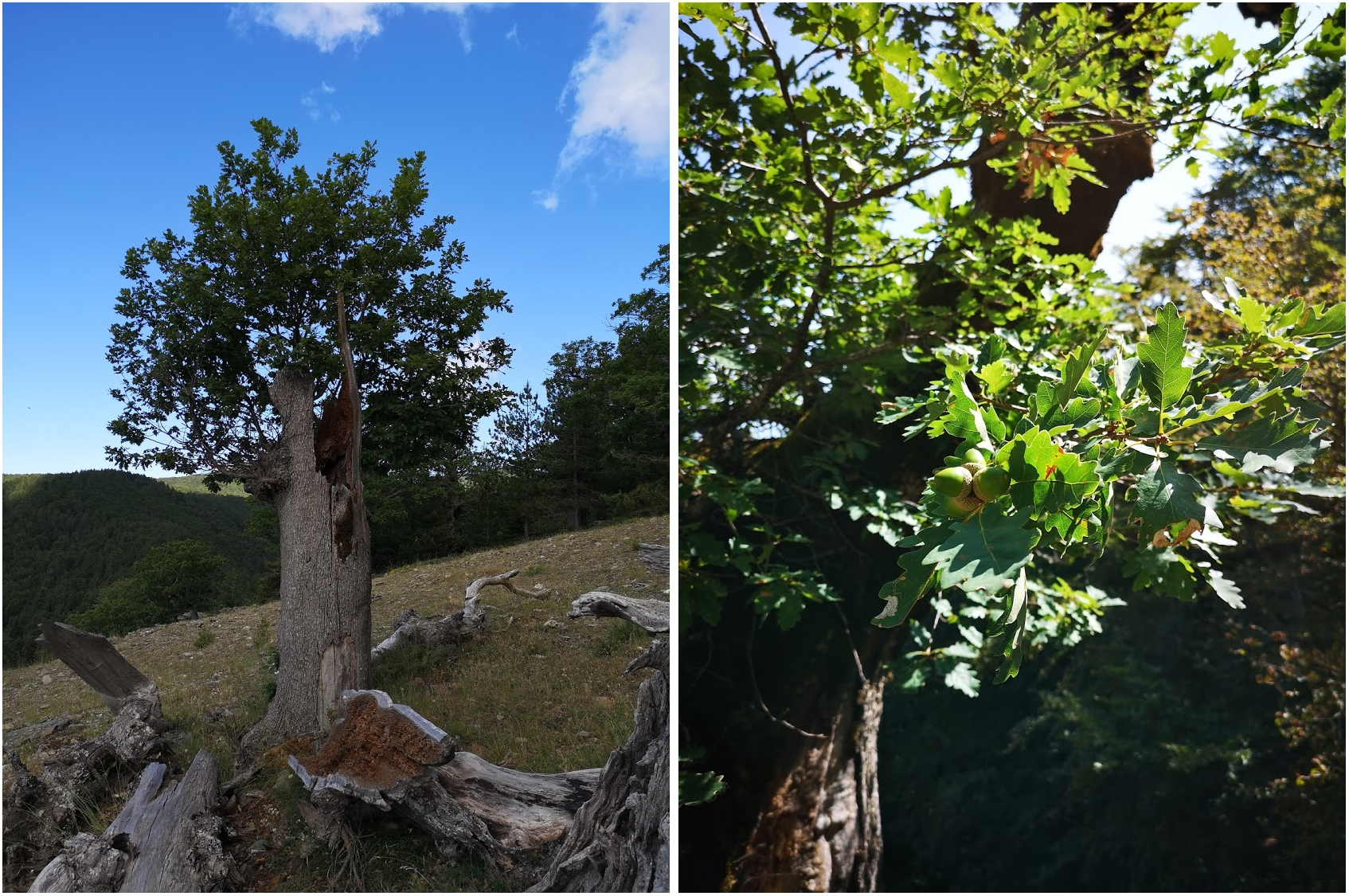
Canopy rejuvenation: epicormic branching after a severe crown disturbance in an old sessile oak (left side). Acorn production in a 600 yr old sessile oak trees in the Aspromonte National Park where the oldest dated temperate flowering tree in the world (930 yrs) is still fruiting (Piovesan et al. 2020).

In fact, the modular tree growth form, through both SAMs and AMs aboveground and root apical meristem (RAMs) and lateral meristem (LMs) belowground, creates a colony of competing genotypes that vary in their genotypic and phenotypic fitness according to their historical development. The cambial meristem continually renews the vascular system of the tree, responding to local conditions, both temporally and spatially, essentially creating a record of change and resilience in the tree wood. Meristematic tissues, both pluripotent and constantly renewing themselves, enable the tree to essentially be potentially immortal ^25^. This pluripotency is also observed in our ability to propagate scion wood from old trees, which is often rejuvenated to some extent when grafted onto a younger tree, truly enabling the possibility of potential immortality for a particular genotype. Ultimately, the lack of separation between the germline and somatic tissue allows the accumulation of heritable somatic mutations ^31,32^ but within-individual selection at the meristem levels allows sweeping out of deleterious mutations and selection for advantageous mutations below the individual level. Epigenetic imprinting ^33-35^ across their canopies over centuries also allows them to be particularly good at tolerating stress and preventing the wear and tear of aging and passing on advantageous traits and phenotypes.

## Living stores of past selection and future adaptive capacity

Abundant evidence suggests that trees can obtain extreme ages, well over a millennium, due to their unique modular growth form and physiological plasticity and that they regularly achieve ages of many centuries. A neutral stochastic death process, as modeled here, produces population age structures that correspond to those observed in natural populations. While the mature and old trees in the population are consistent across model results, the properties of ancient trees only fully develop in large populations with relatively low mortality rates after many centuries of forest growth. These ancient trees reach ages substantially greater than other trees in the community and comprise a small proportion of the total population. We argue that they play a significant and underappreciated role in the evolutionary dynamics of forest tree species.

Critically, in converted forests or populations of very small size, the long demographic tail of ancient trees is missing. Our results suggest that climax forest containing ancient trees obviously requires several centuries, even when mortality rates are relatively high (3%). Estimates that ‘old-growth’ forest can be achieved in 150 ^36^ neglects the impact of ancient trees. These ancient trees are indicators of the degree of development in old-growth processes ^37^. Old and ancient trees are unique proxies for reconstructing past climate and environment ^38^. Such old trees have survived multiple decadal (AMO and PDO generally lasting between 30-70 years) and even longer contrasting climatic phases (Medieval warm period, Little ice age, global warming). While our model assumes neutrality, the survivorship of some trees also indicates the growth conditions where functional traits ^39^ confer resistance to biotic and abiotic stresses. Our currently highly fragmented, largely converted global forests face profound evolutionary challenges, given the loss of the buffering effect of ancient trees, particularly in the face rapidly emerging novel environments expected for the Anthropocene. Once disturbed by man, a regeneration forest needs a millennium to recover its natural stock of ancient trees.

Finally, ancient trees represent a major reservoir for deep genetic diversity that can bridge long temporal gaps between extreme environmental conditions. While these ancient individuals are relatively infrequent in the population, their existence in the community can have profound impacts on the evolutionary dynamics, particularly regarding coalescent and lineage sorting processes, which are both defined by generation time ^40,41^. Given our poor understanding of demographics or population dynamics in forest trees, the impact of possible variation in generation time, caused by skewed or unusual demographic structures ^42-44^ is not frequently considered. We demonstrate that the presence in ancient trees in the community greatly extends generation time and greatly increases its variance, thus also extending and increasing the variance in coalescent times and lineage sorting processes. While evidence of senescence, through diminished seed output, has been recently reported ^45^, older trees often do contribute a disproportionate number of seeds during a fruiting event ^46^ and no evidence exists in regard to their pollen contribution. Their participation in population genetics of the community remains significant, despite their rarity in the community. Empirically, tree genomes are often noted for their high levels of heterozygosity and typically exhibit higher within populations levels of genetic diversity in comparison to between population levels, supporting this hypothesis. This suspended process of coalescence would also address the paradox of fast microevolution but slow macroevolution ^46^.

For all of these reasons, old-growth forests with their unique stock of ancient trees are becoming increasingly important to protect. Losing these trees is similar to allowing species to go extinct. Ecologically, ancient trees are known to be unique biodiversity hubs ^47^ that provide key or unique ecosystem functions unparalleled by managed forests. They contribute disproportionately to the forest rate of carbon sequestration, as this rate continuously increases with tree size ^48^. But these ancient trees, perhaps most critically, are an irreplaceable evolutionary resource for the tree species themselves. The loss of these ancient trees can greatly reduce the evolutionary potential of the species. Preserving and restoring ancient trees everywhere in the world, from the heart of old growth forests to tiny fragments along roadsides, is an urgent goal for a sustainable future. We strongly advocate research focused on these ancient trees and their contribution to the future adaptive capacity of our global forests.

## Methods

### Simulation model

A simple technique for imposing a stochastic mortality rate on a population of individuals with a constant number of individuals in the population was written in Mathematica 12 for Mac OS (Wolfram Research). The simulation involves no spatial or fitness component into the likelihood of death but instead all individuals are equally likely to perish. The initial settings mimic recovery of a site after a complete conversion with all individuals starting at the same time and same age. All individuals are considered established once they appear in the population simulation with equal probability of mortality.

Each step in the simulation is equivalent to a year and the dead individuals are chosen randomly and then removed from the population, with the number of individuals to be removed being the annual mortality rate multiplied times the population size. Surviving individuals have a year added to their age and a new cohort of one year old individuals replace the dead individuals removed from the population. This step is repeated for the duration of the simulation model. All models were run for an equivalent of 15000 years.

The parameters explored in the model vary by population size (100, 1000, 10K, 100K, and 500K individuals) and annual stochastic mortality rate (0.001, 0.005, 0.01, 0.015, 0.02, 0.03, and 0.05). Each set of parameters were examined across 25 replicates. To explore the impact of increasing population size, one set of replicates were also run for one million individuals given a mortality rate of 0.015. Little variation was observed in the results and further exploration of a broader parameter space did not seem warranted. The mortality rate of 0.02 appears to be a bit of a threshold as the results change quickly below this rate but the lower bound of mortality studied here (0.001) does not appear biological relevant as the results do not reflect empirical evidence from global forests or other organisms which match this mortality model. More detailed exploration of the impact of small population sizes on these dynamics is warranted, as the lower bound of 100 individuals generated unusual results and presented certain challenges to the model effort.

To simplify the analysis of the demographic patterns, samples were taken of the population every 100 years during the 15000 year time period and individuals were grouped by age class. All subsequent analyses were performed on these demographic tables. Given the initial conditions of the model, where an even-aged cohort of individuals dominate the population in the beginning of the model but slowly go through attrition until the youngest age class becomes the most abundant. We determined that this point where the age class structure becomes stable represents the population climax age, at which the impact of the reset no longer has an effect. Population climax age was technically defined as the point at which the oldest age class is represented by a single individual. These dynamics are explained in greater detail in the Supplementary Material 1. All analyses were performed after the demographic patterns became stable.

### Age group classification

We detected an emerging group of individuals in our simulations that were substantially older than the rest of the population. We deemed these individuals as ‘ancient’ and described them as life history lottery winners because they were always a small fraction of the overall population but departed entirely from the population characteristics, extending a very long and thin tail to the distribution of ages (see Supplementary Material 2). Further exploration of the age-rank distribution revealed that three separate segments of the population (mature, old, and ancients) could be identified objectively by age, corresponding to the different parts of the age-rank curve. The threshold between the mature and elder groups could be efficiently determined as twice the age of the median age of the entire population. This threshold, easily calculated, corresponds closely to the inflection point in the rank-age distribution where the steady and linear progression becomes non-linear. The threshold age between old trees and ancients was technically defined as the point where the subset of five consecutive ages present in the population are separated by an average of two years, e.g. the sequence of ages becomes 450, 452, 454, 456, 458. We found that applying this rule to different population sizes resulted in different threshold ages for ancients, that increased with population size, but the absolute number of individuals above this threshold remained constant across population sizes. But, plotting this threshold on the rank-age distributions revealed that it did not correspond well with the properties of the curve and only included a very small portion of the obvious long tail of very old individuals. Instead, we found that the 10K individual population appears to represent a point at which the dynamics become predictable at larger population sizes. Setting the threshold for ancient age as the point where the rule was accepted for 10K for all population sizes then fit the properties of the curve.

### Exploring environmental fitness dynamics

In order to explore the role of individuals in these different age groups given cyclical environmental change, we simulated an environmental fitness value that is the summation of four different cycles that recur over the time period of the simulation. Because of the large possible parameter space given this type of model, we wrote a ‘tunable’ but arbitrary equation that generated an emergent pattern that was appropriate to empirical patterns of environmental cycles that occur on various temporal scales, from decades to millennia. The equation is the summation of

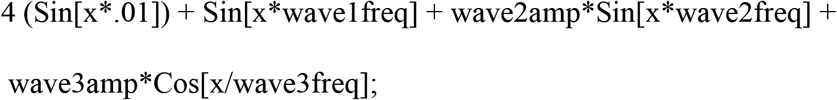

where x = time and the variables wave#amp and wave#freq correspond to the properties of the specific environmental cycle. These variables could be easily modified and a representative set were chosen to examine the impact of the different age groups over these fitness cycles. See Supplementary Materials 3 for a detailed description of the process and the background for the background environmental fitness value used in Figure 3.

To assess the impact of the different age groups on the adaptive capacity of the population, we assumed that fitness determined the ability of the individual to establish. Therefore, the individuals was assigned the fitness value present in the environment at the time of its establishment. For illustrative purposes, a single sample point was chosen arbitrarily for a single replicate (year 10300 in the tenth replicate) to examine the distribution of fitness values for each age group. We feel this approach is adequate and corresponds to our ability to collect empirical data on these types of long-lived individuals, which is inherently latitudinal and not longitudinal in quality. Additionally, this type of analysis is novel and protocols for exploring the properties of population fitness would require the development of a new set of tools. This future work would be required to comprehensively explore the full impact of different age groups across all environmental situations but we feel that this effort is beyond the scope of the current analysis.

The population descriptors of fitness for each age group were compared to the overall background fitness value over 1000 years, across population sizes and mortality rates.

### Estimation of generation time

Generation time was estimated in three ways: 1) all living individuals in a cohort have equal probability of contributing an offspring to the establishing cohort, regardless of their age; 2) individuals in each age class have the same probability of contributing an offspring to the establishing cohort, regardless of their abundance; and 3) only the oldest living individual contributes to the establishing cohort. These three estimators were calculated 1) by randomly choosing an individual from the last living cohort produced at the end of a simulation which served as the ‘tip’ individual; 2) the time of establishment for the tip individual was determined by subtracting the individual’s age from the time of sampling in the simulation; 3) the progenitor of the individual is selected based upon the three criteria described above: i) randomly among all living individuals; ii) randomly among all age classes present; and iii) the oldest living individual. Steps 2 and 3 were repeated until the beginning of the simulation was reached, keeping track of the number of individuals required to traverse the entire simulation, from the end to the beginning. Generation time was then calculated by dividing the length of the simulation (15000 years) by the number of individuals required to span from beginning to end. This entire process was repeated ten times for each of the 25 replicates, meaning that a total of 250 estimates for generation time were calculated for each combination of model parameters.

**Table 1.** Variation in generation time, given different community sizes (rows) and mortality rates (columns). The values for each combination of parameters illustrate mean generation time assuming contribution to the next generation is i) equally among all individuals; ii) proportional to age class; and iii) only by the oldest individual, respectively. The first assumption estimates the average generation time if fecundity is equal across ages and establishment is stochastic. The second assumption estimates the average generation time if fecundity increases linearly with age and establishment is stochastic. The last assumption estimates the maximum possible generation time, given the presence of ancient individuals. Because the temporal resolution of the analysis is in one hundred year steps, the lowest value for generation time is 100 years. The value for one million individuals at 1.5% mortality is 299.

|  | 1.0% |  |  | 1.5% |  |  | 2.0% |  |  | 5.0% |  |  |
| --- | --- | --- | --- | --- | --- | --- | --- | --- | --- | --- | --- | --- |
|  | i | ii | iii | i | ii | iii | i | ii | iii | i | ii | iii |
| 100 | 121 | 121 | 449 | 102 | 103 | 206 | 102 | 103 | 207 | <100 | <100 | <100 |
| 1000 | 121 | 159 | 649 | 106 | 125 | 432 | 102 | 111 | 319 | <100 | <100 | 105 |
| 10000 | 121 | 239 | 822 | 106 | 174 | 563 | 102 | 140 | 415 | <100 | 100 | 136 |
| 100000 | 121 | 341 | 992 | 106 | 232 | 689 | 102 | 184 | 515 | <100 | 103 | 198 |
| 500000 | 121 | 403 | 1121 | 106 | 281 | 776 | 102 | 218 | 588 | <100 | 108 | 213 |

**Supplementary Material 1.**
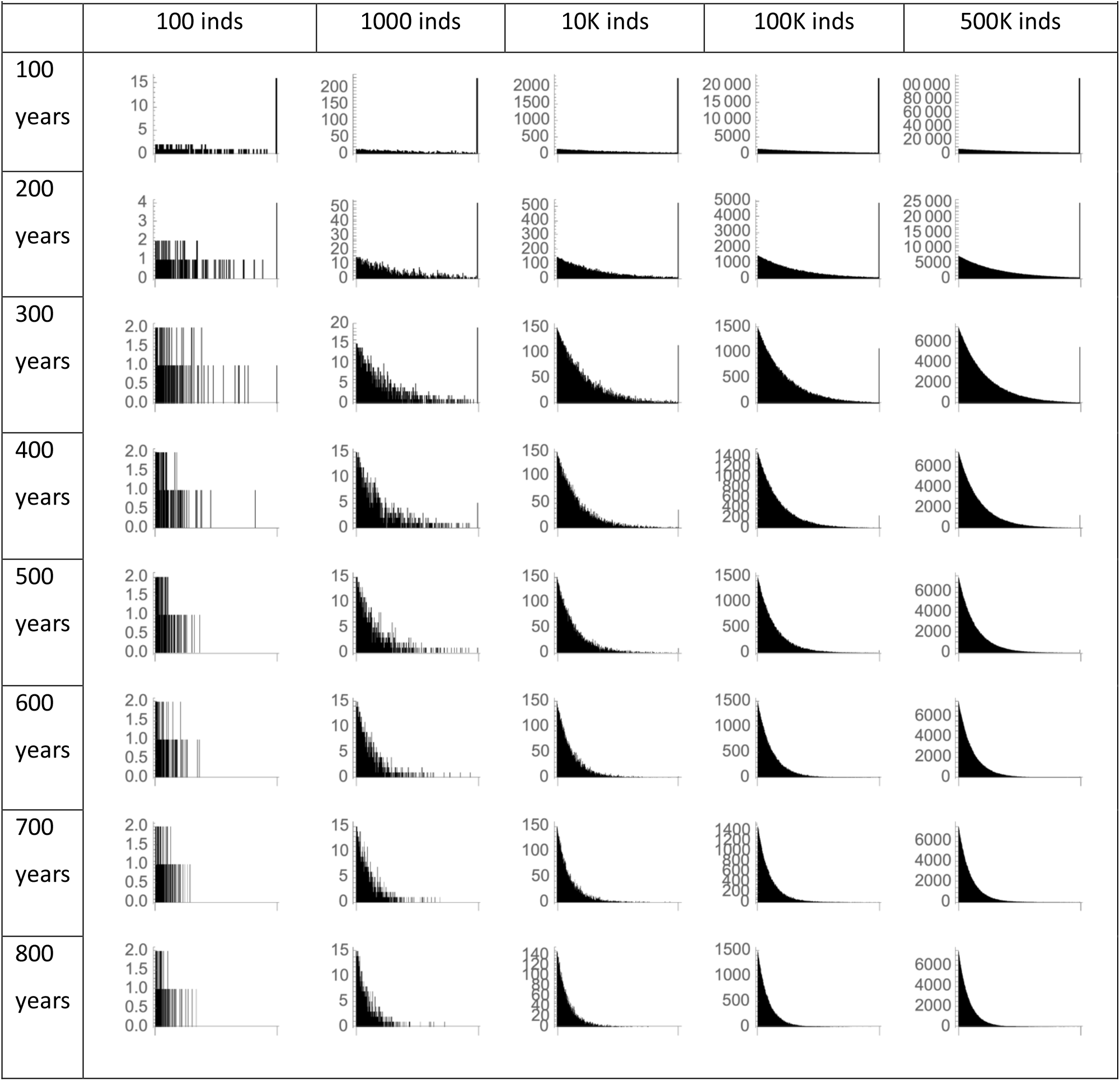
Abundance of age classes in populations of different sizes with increasing population age, in 100 year increments, given an annual mortality rate of 1%. Columns show different population sizes and rows show increasing community age. In each figure, age classes are shown on the x-axis with a maximum value that equals the community age while abundance of individuals in each age class are shown on the y-axis. Values shown are from a single representative replicate.

**Supplementary Material 2.**
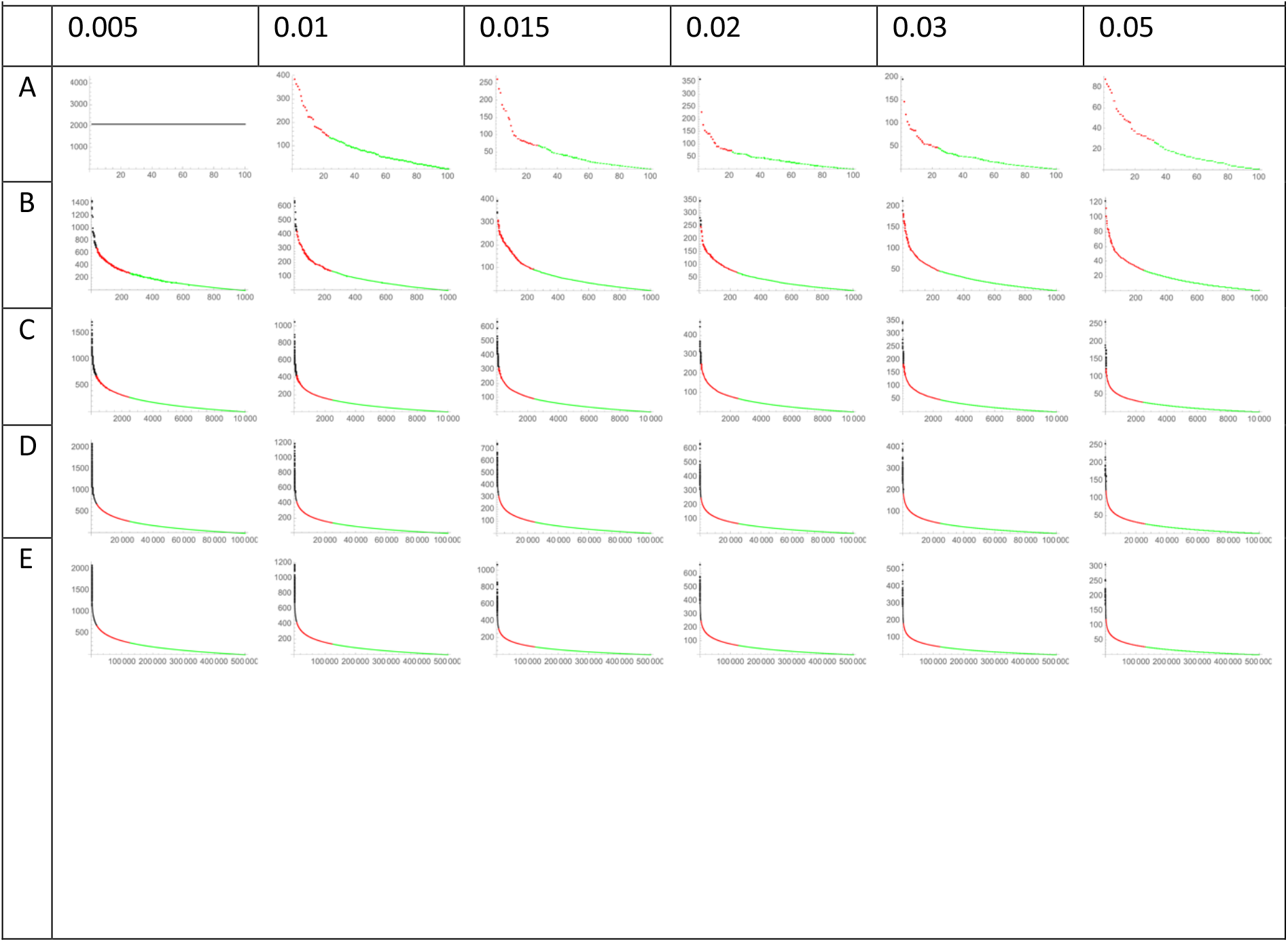
Age-rank distributions of different population sizes and mortality rates. [[what is the time point??]] Population size is shown in rows: A) 100; B) 1000; C) 10K; D) 100K; E) 500K. Mortality rates are shown in the columns. Individuals are arranged according to age, shown on the y-axis, from oldest to youngest, left to right. Individuals are color-coded: ancient = black,old trees= red, young = green.

**Supplementary Material 3.**
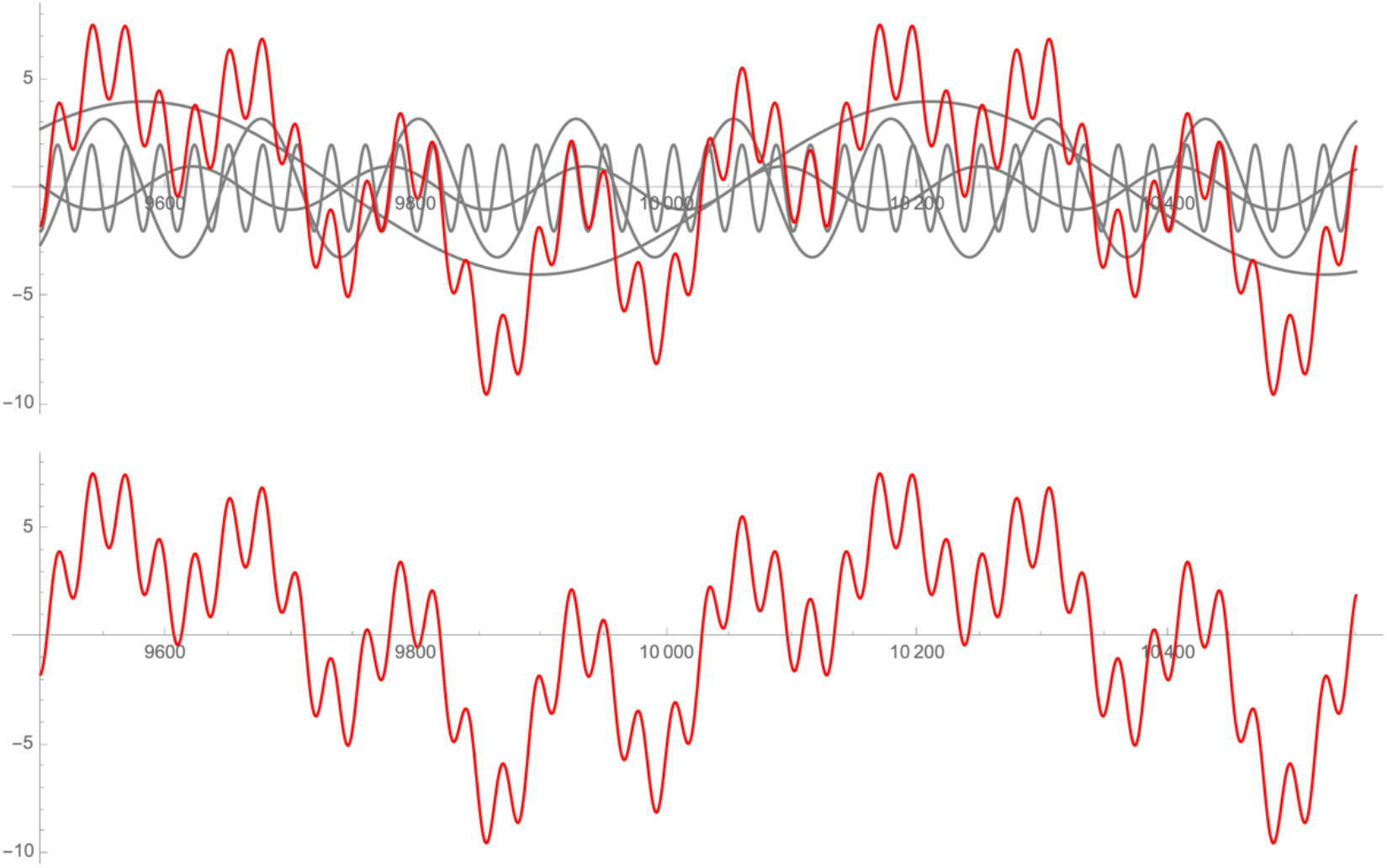
Exploring population level fitness, given multi-cyclical environmental change. Environmental change was modeled by adding four environmental cycles to create an emergent environmental cycle. The top panel shows the four underlying cycles in gray and emergent cycle in red. The bottom panel shows just the emergent cycle.

The functions describing the four underlying cycles are: 1) Sin[x*.01]); 2) Sin[x*wave1freq];

3) wave2amp*Sin[x*wave2freq]; and 4) wave3amp*Cos[x/wave3freq, where x = time point in simulation and wave#amp and wave#freq are variables describing the amplitude and frequency of the cycle.

Population fitness was estimated within a one thousand year time period to reflect a reasonable amount of environmental change during which each underlying cycle and the emergent cycle would go through several complete revolutions.

---

**Supplementary Material.**
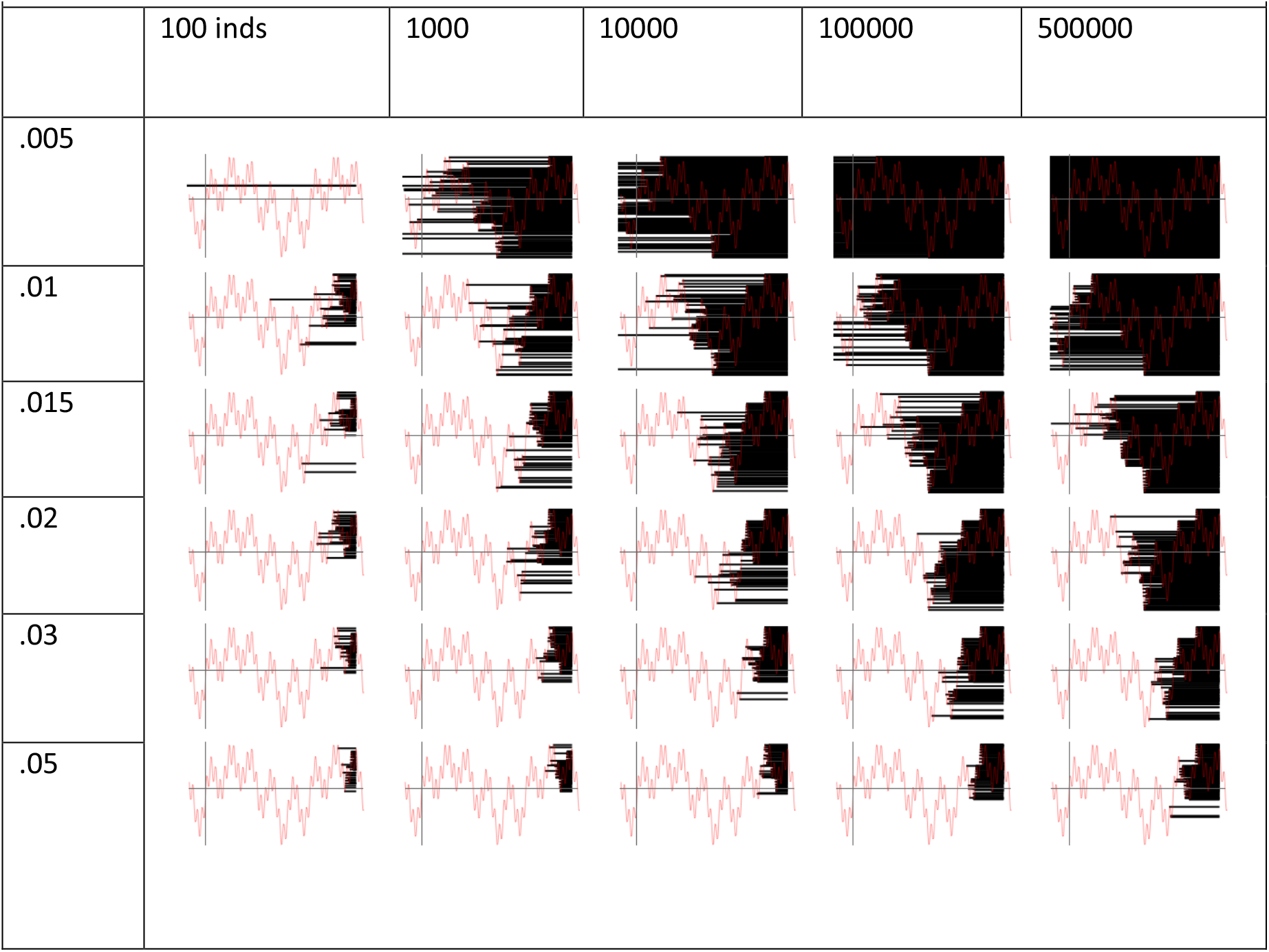
Demographic patterns of populations given different mortality rates (rows) and population sizes (columns). The red line indicates the fluctuating fitness value determined by four interacting environmental cycles. The black lines represent the individuals alive at the end of the time series, originating at the time of their establishment, for each set of parameter conditions. This illustration represents one replicate and one time point.

